# Improving the accuracy of two-sample summary data Mendelian randomization: moving beyond the NOME assumption

**DOI:** 10.1101/159442

**Authors:** Jack Bowden, Fabiola Del Greco M, Cosetta Minelli, Qingyuan Zhao, Debbie A Lawlor, Nuala A Sheehan, John Thompson, George Davey Smith

## Abstract

**Background:** Two-sample summary data Mendelian randomization (MR) incorporating multiple genetic variants within a meta-analysis framework is a popular technique for assessing causality in epidemiology. If all genetic variants satisfy the instrumental variable (IV) and necessary modelling assumptions, then their individual ratio estimates of causal effect should be homogeneous. Observed heterogeneity signals that one or more of these assumptions could have been violated.

**Methods:** Causal estimation and heterogeneity assessment in MR requires an approximation for the variance, or equivalently the inverse-variance weight, of each ratio estimate. We show that the most popular ‘1st order’ weights can lead to an inflation in the chances of detecting heterogeneity when in fact it is not present. Conversely, ostensibly more accurate ‘2nd order’ weights can dramatically increase the chances of failing to detect heterogeneity, when it is truly present. We derive modified weights to mitigate both of these adverse effects.

**Results:** Using Monte Carlo simulations, we show that the modified weights outperform 1st and 2nd order weights in terms of heterogeneity quantification. Modified weights are also shown to remove the phenomenon of regression dilution bias in MR estimates obtained from weak instruments, unlike those obtained using 1st and 2nd order weights. However, with small numbers of weak instruments, this comes at the cost of a reduction in estimate precision and power to detect a causal effect compared to 1st order weighting. Moreover, 1st order weights always furnish unbiased estimates and preserve the type I error rate under the causal null. We illustrate the utility of the new method using data from a recent two-sample summary data MR analysis to assess the causal role of systolic blood pressure on coronary heart disease risk.

**Conclusions:** We propose the use of modified weights within two-sample summary data MR studies for accurately quantifying heterogeneity and detecting outliers in the presence of weak instruments. Modified weights also have an important role to play in terms of causal estimation (in tandem with 1st order weights) but further research is required to understand their strengths and weaknesses in specific settings.

## Introduction

Mendelian randomization (MR) [1] is an instrumental variable approach that uses genetic data, typically in the form of single nucleotide polymorphisms (SNPs), to assess whether a modifiable exposure exerts a causal effect on a health outcome in the presence of unmeasured confounding. A particular MR study design gaining in popularity instead combines publically available summary data on SNP-exposure and SNP-outcome associations from two separate studies for large numbers of uncorrelated variants within the framework of a meta-analysis. These studies should contain no overlapping individuals (to ensure independence) but should also originate from the same source population. This is referred to as two-sample summary data MR [2]. Providing the necessary modelling assumptions are met and the chosen set of SNPs are all valid instrumental variables, an inverse-variance weighted (IVW) average of their individual causal ratio estimates provides an efficient and consistent estimate for the causal effect. This is referred to as the IVW estimate (see **Box 1**). Cochran’s *Q* statistic, which is derived from the IVW estimate, should follow a *χ*^2^ distribution with degrees of freedom equal to the number of SNPs minus 1. Excessive heterogeneity is an indication that either the modelling assumptions have been violated, or that some of the genetic variants violate the IV assumptions - for example, by exerting a direct effect on the outcome not through the exposure [3]. This is termed ‘horizontal pleiotropy’ [4, 5]. For brevity we will refer to problematic horizontal pleiotropy simply as pleiotropy from now on.

### Box 1: Standard two-sample summary data MR

**The IV assumptions:** The canonical approach to MR assumes that group of SNPs are valid IVs for the purposes of inferring the causal effect of an exposure, *X*, on an outcome, *Y*. That is they are: associated with *X* (IV1); not associated with any confounders of *X* and *Y* (IV2); and can only be associated with *Y* through *X* (IV3). The IV assumptions are represented by the solid lines in the causal diagram below for a SNP *G*_*j*_, with unobserved confounding represented by *U*. Dotted lines represent dependencies between *G* and *U*, and *G* and *Y* that are prohibited by the IV assumptions. The causal effect of a unit increase in *X* on the outcome *Y*, denoted by *β*, is the quantity we are aiming to estimate.

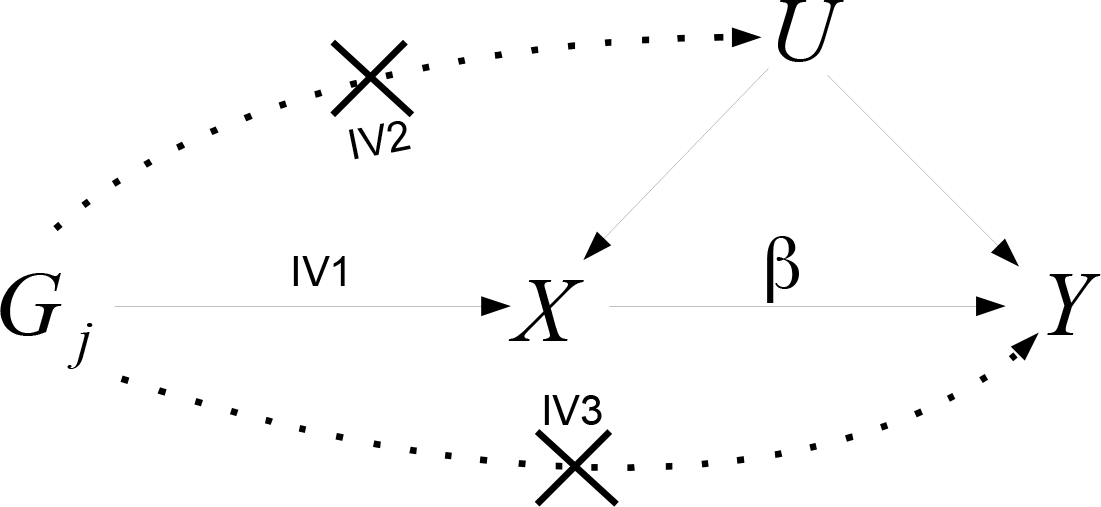

**The Ratio estimate:** Assume that exposure *X* causally affects outcome *Y* linearly across all values of *X*, so that a hypothetical intervention which induced a 1 unit increase in *X* would induce a *β* increase in *Y*. Suppose also that all *L* SNPs predict the exposure via an additive linear model with no interactions. If SNP *j* is a valid IV, and the two study samples are homogeneous, then the underlying SNP-outcome association from sample 1, Γ_*j*_, should be a scalar multiple of the underlying SNP-exposure association estimate from sample 2, *γ*_*j*_, the scalar multiple being the causal effect *β*. That is:

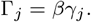

The ratio estimate for the causal effect of *X* on *Y* using SNP *j* (out of *L*), 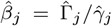, where 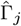 is the estimate for SNP *j*’s association with the outcome (with standard error *σ*_*Y j*_) and 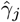 is the estimate for SNP *j*’s association with the exposure (with standard error *σ*_*X j*_).

**The IVW estimate:** The overall inverse-variance weighted (IVW) estimate for the causal effect obtained across *L* uncorrelated SNPs is then given by

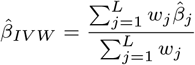

where *w*_*j*_ is the inverse-variance of 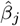 Cochran’s *Q* statistic:

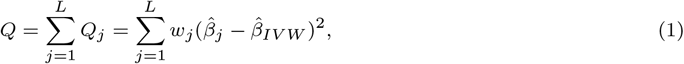

can then be used to test for the presence of heterogeneity. If heterogeneity is detected, this provides evidence of horizontal pleiotropy. Two popular choices for the inverse-variance weights used to calculate the IVW estimate and Cochran’s *Q* statistic are:

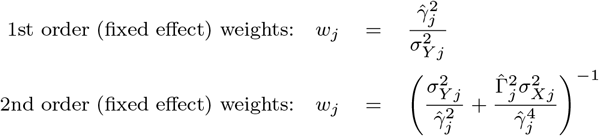

In the two-sample setting, 2nd order weights are simplified because it is not necessary to include terms involving the covariance of 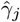 and 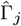, since they are obtained from independent samples. For a more detailed description of the assumptions required by two-sample summary data MR, see Bowden et al. [7].

The presence of heterogeneity due to pleiotropy does not necessarily invalidate an MR study. If across all variants: (i) the amount of pleiotropy is independent of instrument strength (the InSIDE assumption [6]) and (ii) it has a zero mean, then a standard random effects meta-analysis will still yield reliable inferences [6, 7]. Although many MR methods now exist which offer robustness to pleiotropy, in this paper we focus solely on the standard IVW estimate.

### Choice of weights in two-sample summary data MR

Typically, ‘1st order’ inverse-variance weights are used to calculate both the IVW estimate and Cochran’s *Q*. 1st order weights ignore uncertainty in the denominator of the ratio estimate, which is equivalent to making the ‘NO Measurement Error’ (NOME) assumption, as defined in [7, 8]. This nomenclature is chosen to remind the practitioner that the SNP-exposure association estimates are only equal to the true associations when measured with infinite precision (or without error). The NOME assumption does not relate to absence of measurement error in the exposure itself, which can also be problematic for MR studies [9]. Although the NOME assumption is never completely satisfied, strong violation (via the use of weak genetic instruments) induces classical regression dilution bias in the IVW estimate towards the null. So called ‘2nd order’ weights, attempt to better acknowledge the full uncertainty in the ratio estimate of causal effect from each SNP [10, 11] (see **Box 1**). It may appear obvious that 2nd order weights should be used as standard within an MR study to calculate the IVW estimate and Cochran’s *Q*. In fact, Thompson et al. [12] showed that 2nd order weighting produces causal estimates which are generally more biased than 1st order weighting. The ability of 1st and 2nd order weighting to furnish reliable *Q* statistics has yet to be fully explored.

## Methods

It is possible to view Cochran’s *Q* statistic not just as a method for quantifying heterogeneity, but as a tool for directly estimating the causal effect. That is, the IVW estimate actually minimises Cochran’s *Q*. We use this fact to derive a generalised estimating equation based on an extended version of Cochran’s *Q* statistic (see **Box 2**), where its weight term is allowed to be a function of the causal effect parameter. We show that 1st order and 2nd order weighting are special cases of this general weight function. Using this formulation we propose two new procedures for causal effect estimation and heterogeneity quantification.

### Box 2: Accounting for weak instruments under a fixed effect model and testing for pleiotropy

We start by writing down two models: firstly the underlying data generating model for the SNP-outcome association estimates under the assumption of no pleiotropy, which is a function of the causal effect and the true SNP-exposure association; and secondly the model that we actually *fit* to the data, which is a function of the causal effect and the SNP-exposure association *estimate*:

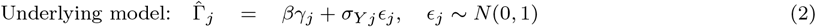

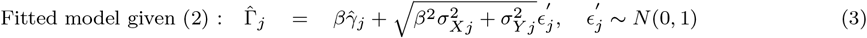

Note that the variance of the error term in the fitted model has been inflated by a factor of 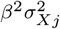 by virtue of replacing *γ_j_* with its estimate in (3). Dividing both sides of the fitted model through by 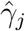 we can obtain a model for the *j*th ratio estimate, and from that an expression for its variance:

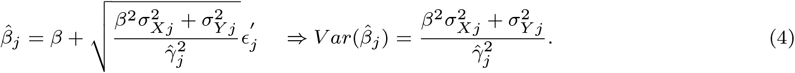

The variance term 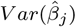 in (4) is a function of the true causal effect *β*. Let its reciprocal inverse-variance weight be denoted as 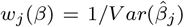. Using this weight, we now define the following modfied *Q* statistic and IVW estimate:

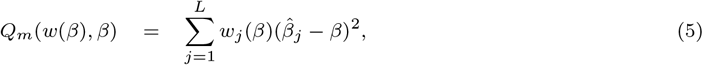

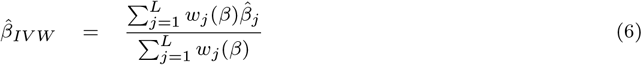

The IVW estimate using 1st order weights is obtained by replacing *w*_*j*_(*β*) with *w*_*j*_(0) in (6). Likewise, its associated heterogeneity statistic is *Q*_*m*_(*w*(0), *β*). The IVW estimate using 2nd order weights is obtained by replacing *w*_*j*_(*β*) with 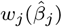 in (6). Likewise, its associated heterogeneity statistic is 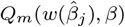.

We now introduce two new fixed effect IVW estimates (and associated heterogeneity statistics) obtained via different weighting schemes.

**The ‘iterative’ IVW estimate** Briefly, let 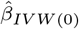 be the IVW estimate obtained using 1st order weights. Now define 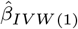 as the IVW estimate obtained from plugging 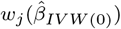 into (6). Lastly, define 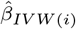 as the IVW estimate obtained from plugging 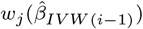 into (6). We call 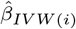 the *i*th ‘iterative’ IVW estimate and 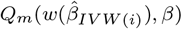 its associated heterogeneity statistic.

**The ‘exact’ IVW estimate** Although we obtain the 1st order, 2nd order and iterative IVW estimates directly from formula (6), each one has the property that it minimises its equivalent *Q* statistic in (5). Crucially, the weights of these *Q* statistics do not depend on *β* because a value (or estimate) has been substituted in its place.

In contrast, the exact IVW estimate is the value obtained from directly minimising the generalised *Q* statistic *Q*_*m*_(*w*(*β*), *β*) in equation (5) with respect to *β*. Here, the weights are now allowed to be a proper function of *β* and affect the minimisation. Letting 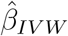 now represent the exact IVW estimate derived in this manner, 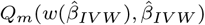 is then its associated heterogeneity statistic.

Our first procedure is termed the ‘iterative’ approach. It iteratively updates the weight term with improved guesses for the causal parameter, using the 1st order IVW estimate as a starting point. This procedure is closely related to the ‘two-step GMM’ estimator [13] used in econometrics. Our contribution has been to describe how it can be implemented using Cochran’s *Q* statistic in the two-sample summary data MR setting. It will be shown that the iterative IVW approach improves causal effect estimation and heterogeneity detection compared to 1st and 2nd order weighting. However, regardless of the number of iterations performed, this procedure will not in general yield the same results as that obtained from directly minimising a more general version of Cochran’s *Q*, where in addition the weight term is allowed to be a proper function of the causal effect parameter *β*. We refer to this second procedure as the ‘exact’ approach. The exact IVW estimate can be viewed as analogous to the limited information maximum likelihood (LIML) estimate, translated to the two-sample summary data MR setting [14]. For further details see **Box 2**.

### Estimation and inference after detection of pleiotropy

**Box 2**describes how to use *Q* statistics to calculate the IVW estimate under a fixed effect model and to test for the presence of heterogeneity due to pleiotropy. If substantial heterogeneity is detected, inferences about the causal effect need to be adjusted to take this additional uncertainty into account, by assuming a random effects model [15, 16]. In Appendix 1 of *Online Supplementary Material* we describe in detail how to generalise the *Q* statistics to obtain point estimates, standard errors and confidence intervals for the 1st order, 2nd order, iterative and exact IVW estimate under both fixed and random effects models (the multiplicative model is currently preferred for MR studies). This task is straightforward for the 1st order, 2nd order and iterative weighting approaches because they can be fitted using standard regression software. Bespoke methods are needed for exact weighting, however, and a short summary of this particular approach is provided in **Box 3**. Specifically, in the fixed effect case we describe how to invert the exact *Q* statistic to get a 95% confidence interval for the exact weighted IVW estimate. In the random effects case, we describe how to jointly estimate the causal effect and multiplicative over-dispersion parameter using a system of two estimating equations. A non-parametric bootstrap algorithm is then proposed to obtain a confidence interval for the causal effect.

#### Box 3: Accounting for weak and pleiotropic instruments using exact weighting

First define the following generalized *Q* statistic and weight function for the multiplicative random effects model:

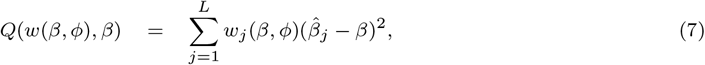

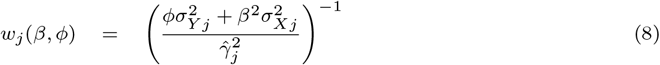

Here *ϕ* (which is greater than or equal to 1) is the multiplicative scale factor that quantifies the degree of heterogeneity.

**Inference for exact weighting under a fixed effect model** When *ϕ* is set to 1 in equations (7) and (8) this is equivalent to assuming a fixed effect model, and minimising (7) with respect to *β* gives the fixed effect exact IVW estimate, as described in **Box 2**. We explore two ways to calculate the standard error of the fixed exact IVW estimate, denoted by 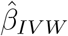. The first method uses the standard error formula:

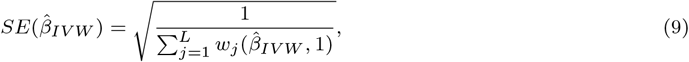

to construct symmetric ninety-five percent confidence intervals for the causal effect as 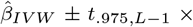 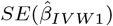. Here *t*_.975,*L*−1_ is the 97.5th percentile of student’s t-distribution with *L* − 1 degrees of freedom. This same procedure is used to derive confidence intervals for the IVW estimate under 1st order, 2nd order and iterative weighting.

The second method directly inverts the *Q* statistic to find the confidence set:

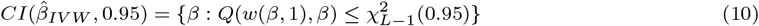

Where 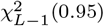 is the 95th percentile of a chi-squared distribution with *L* − 1 degrees of freedom. In order to improve the properties of this approach with few instruments, we additionally replace the value 0.95 in (10) with the value 2Φ(*z*) − 1, where *z* is the 97.5th percentile of a *t*-distribution with *L* − 1 degrees of freedom and Φ() is the cumulative distribution function of a standard normal distribution. As *L* increases, 2Φ(*z*) − 1 tends to 0.95 from above.

**Inference for exact weighting under a random effects model** The fixed effect exact IVW estimate, and its associated confidence intervals will only give reliable estimates if the fixed effect model holds. In practice, substantial heterogeneity is generally present in MR studies, in which case a random effects model should be adopted. The random effects exact IVW estimate is obtained by finding the joint value of (*β*,*ϕ*) that solves:

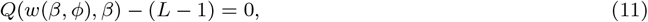

subject to the constraint that

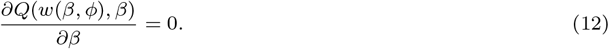

It is not straightforward to obtain a reliable confidence interval for the causal parameter *β* using the inversion method - as in equation 10 - when over-dispersion is allowed. This is because it ignores uncertainty in the estimation of *ϕ*. Instead we obtain an estimate for the variance of 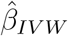 using a standard non-parametric bootstrap algorithm. For further details please see Appendix 1 of *Online Supplementary Material*.

### Performance of the *Q* statistics under no pleiotropy

We now assess the extent to which *Q* statistics derived using 1st order, 2nd order, iterative and exact weighting erroneously detect heterogeneity due to pleiotropy when it is not present (i.e. its type I error rate). To assess this, two-sample summary data MR studies comprising 25 SNP-exposure and SNP outcome association estimates were generated from models with no heterogeneity due to pleiotropy. This furnished a set of ratio estimates between which no additional variation should exist as their instrument strength grows large (because NOME is satisfied), or if the causal effect (*β*) equals zero. To highlight this we simulated MR studies with a range of instrument strengths - from weak (a mean *F*-statistic of 10) to strong (a mean *F*-statistic of 100). Further details of the simulation study set up are described in Appendix 2 of *Online Supplementary Material*.

Table 1 (columns 2-9) show the mean *Q* statistic and the probability of the *Q* statistic detecting heterogeneity at the 5% significance level (the type I error rate), when using 1st order, 2nd order, iterative and exact weights. Five different mean *F*-statistic values were considered for *β*=0 (no causal effect), *β*=0.05 and *β*=0.1, giving 15 scenarios in total. Four iterations were used for the iterative weighting method, as this was sufficient to ensure convergence. We note that in the absence of a causal effect (*β*=0), 1st order weights are exactly correct. Furthermore, in the presence of a causal effect, when the mean *F*-statistic is 100 all weighting methods are near-exact. Under the causal null, all weighting schemes control the type I error rate for detecting heterogeneity. 2nd order weighting is extremely conservative in this respect with weak instruments, however (e.g. a type I error rate near zero when *F*=10).

**Table 1:**
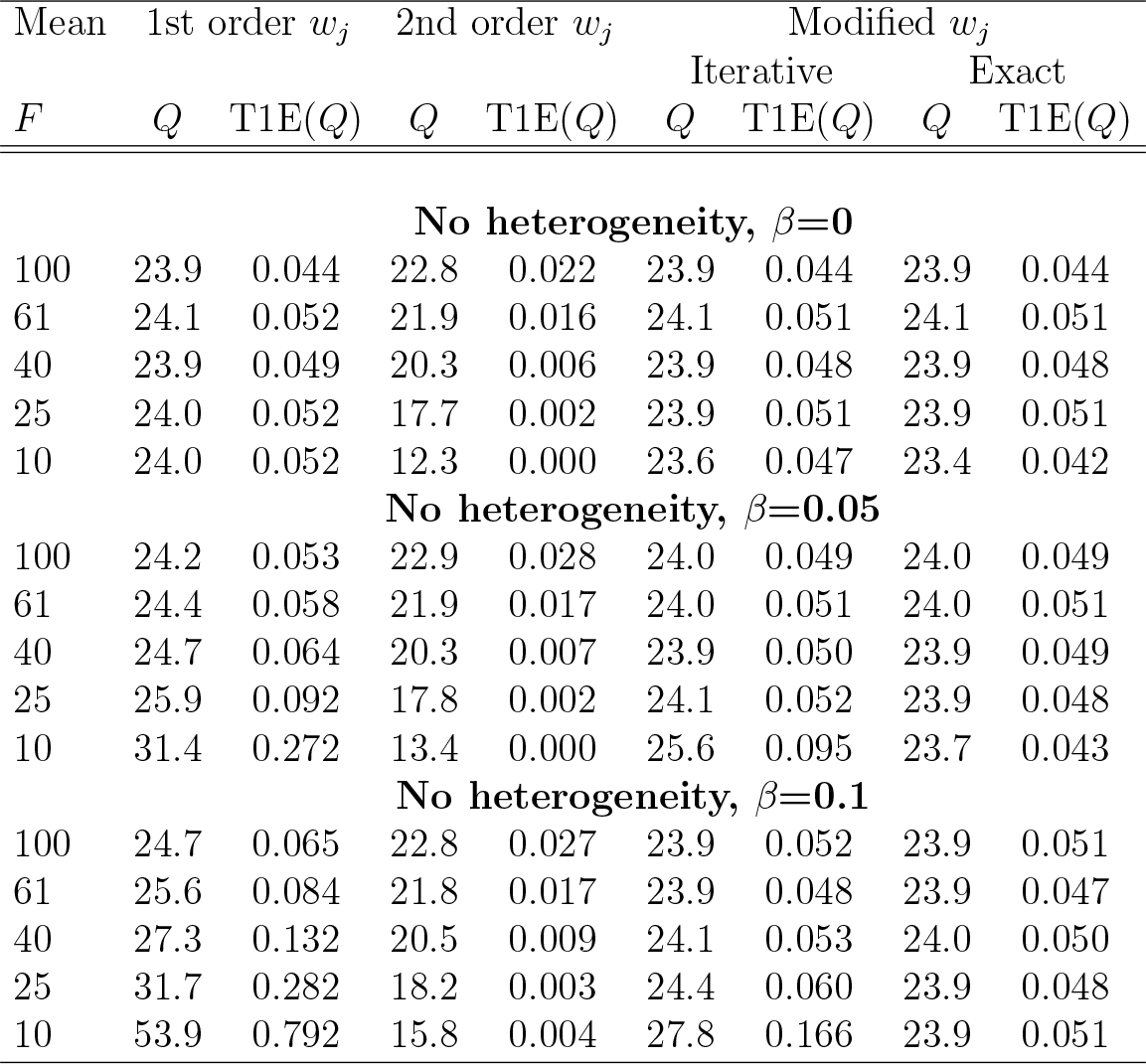
Mean Q statistic and type I error rate (T1E) of 1st order, 2nd order, iterative (four iterations were performed) and exact weighting. Results are the average of 10,000 simulated data sets. Type I error rate (T1E(Q)) refers to the proportion of times Q is greater than the upper 95th percentile of a 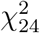 distribution.

In the presence of a causal effect, 1st order weights under-estimate the true variability amongst the ratio estimates as the mean *F*-statistic reduces. The associated *Q* statistics are then too large on average (i.e. positively biased beyond their expected value of 24). This inflates the type I error rate for detecting pleiotropy beyond nominal levels (e.g. a type I error rate of ≈ 80% when *F*=10 and *β*=0.1). 2nd order weighting continues to over-correct the *Q* statistic so that it is negatively biased, thereby removing *any* ability to detect heterogeneity at all. In contrast, iterative weights are much more effective at preserving the type I error rate of the *Q* statistic at its nominal level, unless the mean *F*-statistic is very low (indicating weak instruments). Exact weighting perfectly controls the type I error rate of Cochran’s *Q* across all the scenarios considered. Appendix 2 of *Online Supplementary Material* shows equivalent results for MR studies of 10 and 100 variants, with highly similar results.

Figure 1 (left and right) shows the distribution of *Q* statistics using 1st order, 2nd order and exact weights for *β*=0.1 and when the mean *F*-statistic is 100 and 10. This illustrates how exact weighting ensures Cochran’s *Q* statistic is faithful to its correct null distribution.

**Figure 1:**
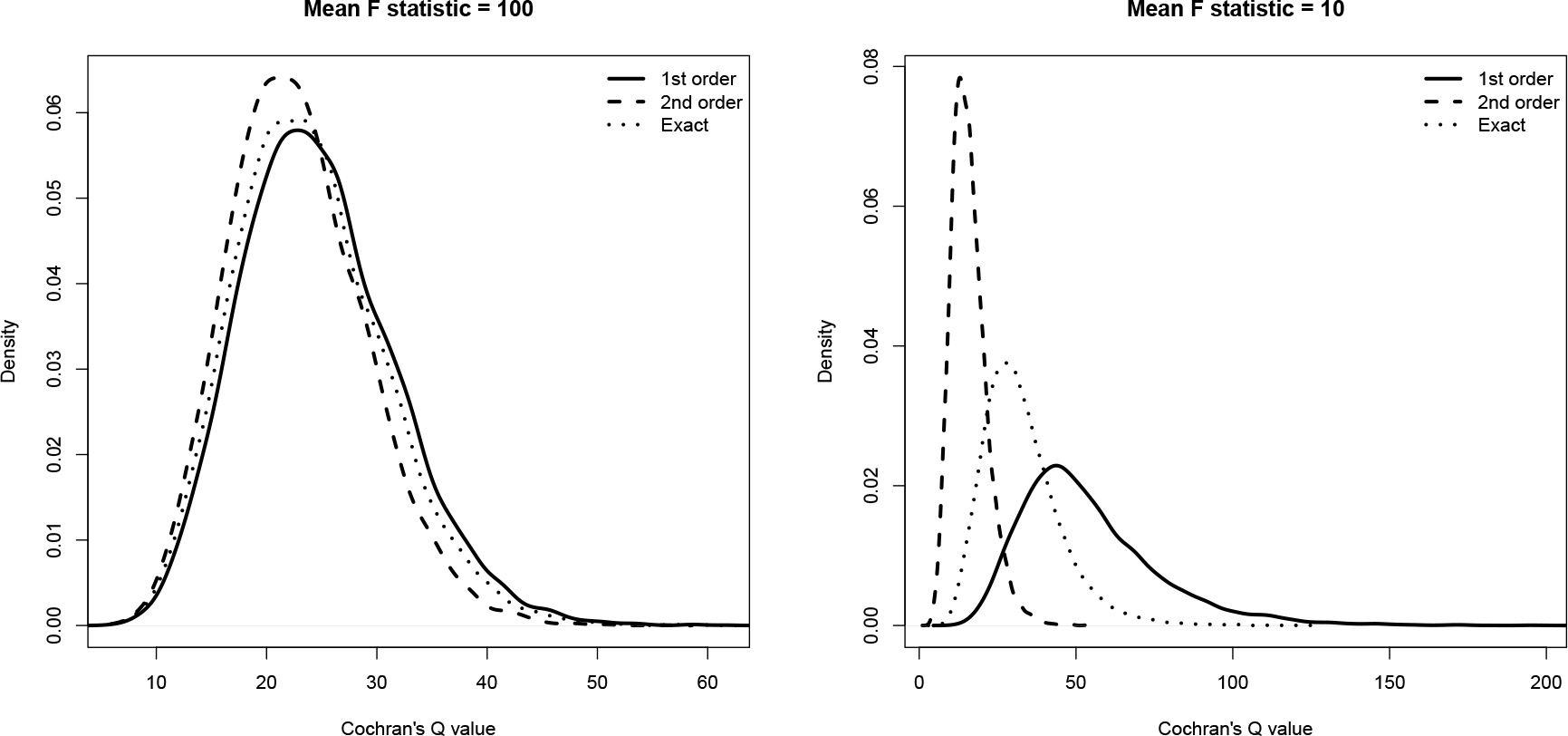
Distribution of Q statistics (with 25 degrees of freedom) using 1st order, 2nd order and exact weights. The causal effect β=0.1 and the mean F statistic equals 100 (left) and 10 (right) respectively.

#### Power to detect pleiotropy

In Table 1 the type I error rate of Cochran’s *Q* statistic for detecting heterogeneity using 2nd order weights was below its nominal level. This is detrimental if it translates into a low statistical power to detect heterogeneity when it *is* truly present. Figure 2 (left) shows the power of Cochran’s *Q* to detect heterogeneity at the 5% significance level as a function of 1st order, 2nd order, iterative and exact weights when data are simulated under a multiplicative random effects model with heterogeneity due to pleiotropy of increasing magnitude (specifically, equation (2) in Appendix 1 of *Online Supplementary Material* was used).

**Figure 2:**
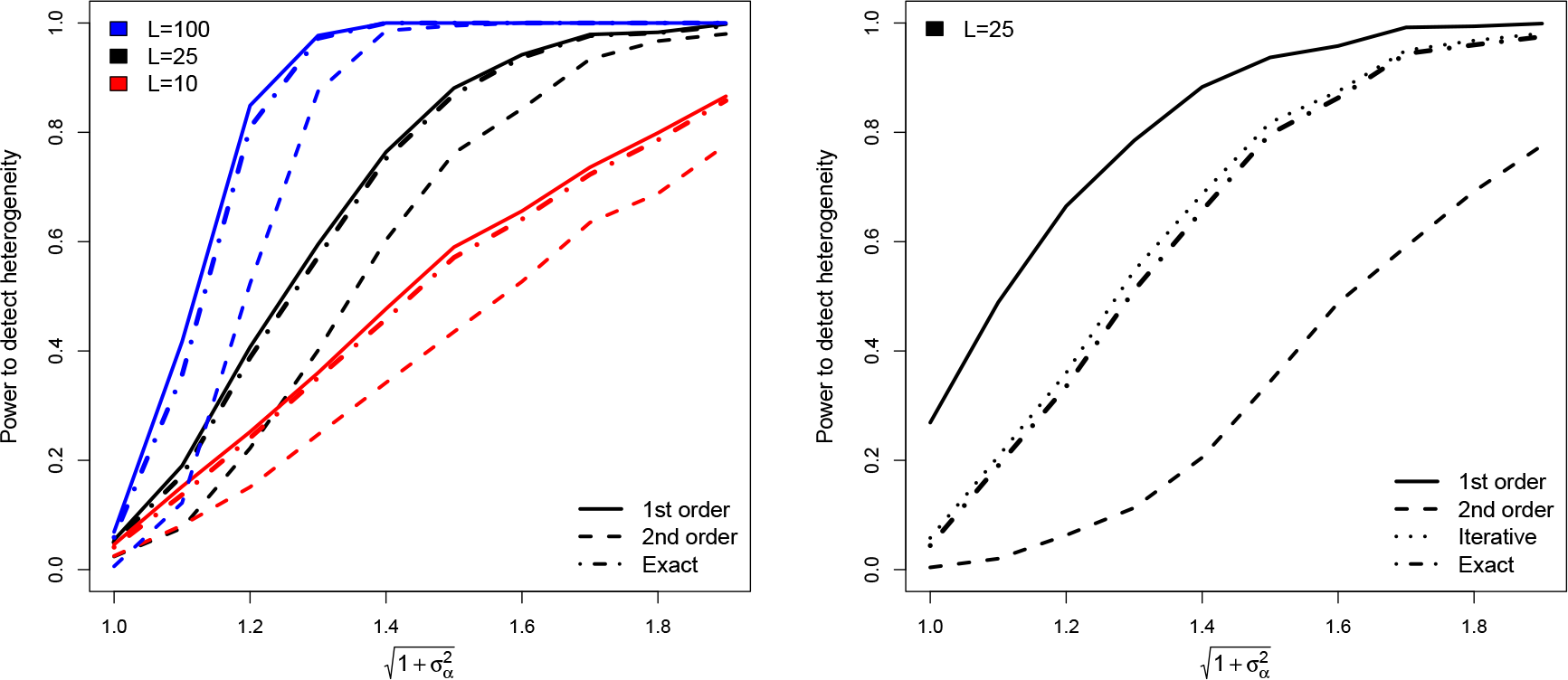
Left: Power of Cochran’s Q statistic to detect heterogeneity as a function of the pleiotropy variance and number of SNPs (L) using 1st order, 2nd order and exact weights. Pleiotropy is simulated under a multiplicative random effects model. The causal effect is equal to 0.05 and the mean F-statistic is 61. Right: Equivalent power plot except the causal effect is equal to 0.1 and the mean F-statistic is 25.

The simulation is repeated for MR analyses with 10, 25 and 100 SNPs. For all simulations, the causal effect equalled 0.05 and the mean *F*-statistic equalled 61. We see that the power of Cochran’s *Q* to detect heterogeneity approaches 100% for all weighting schemes as the pleiotropy variance increases. Power also increases with the number of SNPs. The power of iterative or exact weights is near identical, therefore we only show results for the exact weights for clarity. The most striking result in this plot is that the power of 2nd order weighting always lags considerably behind that of 1st order or exact weights.

Figure 2 (right) shows the results of a near identical simulation for the case *L*=25, except that the causal effect is set to 0.1 and the mean *F*-statistic is equal to 25. We see that the power to detect heterogeneity is always greatest when using 1st order weights, but only because its power curve starts at a baseline level of 28% when there is no pleiotropy. This corresponds to the type I error rate observed in row 14 of Table 1. The power of iterative and exact weighting starts at the correct 5% level, and rapidly increases to 100% as the pleiotropy variance increases. The two implementations of our modified weights can be differentiated in this simulation, with the iterative approach being slightly more powerful. The power of 2nd order weighting, unsurprisingly, lags considerably behind the rest. Equivalent plots for data simulated under an additive pleiotropy model are shown in Appendix 3 of *Online Supplementary Material*, and are highly similar.

#### Detecting outliers using individual components of *Q*

When heterogeneity is detected by the IVW model, it is interesting to investigate whether this is contributed to by all SNPs, or if instead a small number of SNPs are responsible. Under the null hypothesis of no heterogeneity, *Q* should follow an appropriate 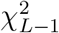 distribution, with *L* being the number of SNPs. Likewise, each individual component of *Q* can be approximated by a 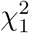 distribution. If an individual SNP’s *Q* contribution is extreme (for example above the 5% threshold of 3.84 or instead a bonferroni corrected threshold), then it may be desirable to exclude the SNP in a sensitivity analysis. Although we do not want to advocate a rigid, blanket policy of outlier removal, in Appendix 4 of *Online Supplementary Material* we illustrate via simulation how the reliability of such a procedure depends on the choice of weights. The simulation (with 26 SNPs and a single larger outlier) is motivated by the real data example in the following section. In this instance, our simulation suggests that iterative rather than exact weights are best at correctly identifying outliers due to pleiotropy.

### Estimator performance with and without pleiotropy

Table 2 shows the performance of the 1st order, 2nd order, iterative and exact weigthing in providing accurate point estimates, standard errors and confidence intervals for the causal effect under a fixed effect (no heterogeneity) model for MR analyses of 25 variants. For exact weighting we show the empirical coverage using two different methods: A symmetric 95% confidence interval (labelled ‘CF_1_’) and a 95% confidence interval obtained from inverting its *Q* statistic (labelled ‘CF_2_’), as described in **Box 3**. Importantly, all methods give reliable unbiased estimates with correct coverages under the causal null hypothesis. In the presence of a non-zero causal effect 1st order and 2nd order IVW estimates are increasingly affected by regression dilution bias (and consequently worsening coverage) as the instrument strength decreases. Iterative weights also produce IVW estimates that suffer from regression dilution bias and sub-optimal coverage, but to a lesser extent than 1st or 2nd order weighting. Exact weighting perfectly removes the effect of regression dilution bias (although the precision of the estimate is reduced) and confidence intervals obtained via the inversion method have the correct coverage. Equivalent results for MR studies with 10 and 100 SNPs are shown in Appendix 5 of *Online Supplementary Material*. When only 10 SNPs are available and they are all weak, the coverage of the inverted confidence interval for the exact IVW estimate is slightly conservative (e.g. 96%-98% instead of 95%). As the number of SNPs increases to 100, coverage is very close to the nominal 95% level irrespective of instrument strength.

**Table 2:** *Mean causal estimate 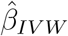, standard error (SE) and coverage frequency (CF) of the 95% confidence interval when using 1st order, 2nd order, iterative and exact weights. Number of variants L=25. CF_1_ = coverage of a symmetric 95% confidence interval, CF_2_ = coverage of inverted Q statistic confidence interval*.

| Mean | 1st order $w_j$ | 2nd order $w_j$ | Modified $w_j$ | | |
| --- | --- | --- | --- | --- | --- |
|  |  |  | Iterative | Exact |  |
| $F$ | $\hat{\beta}_{IVW}(SE); CF$ | $\hat{\beta}_{IVW}(SE); CF$ | $\hat{\beta}_{IVW}(SE); CF$ | $\hat{\beta}_{IVW}(SE); CF_1$ | $CF_2$ |
| **No Heterogeneity, $\beta=0$** |  |  |  |  |  |
| 100 | 0.000 (0.011); 0.952 | 0.000 (0.011); 0.951 | 0.000 (0.011); 0.952 | 0.000 (0.011); 0.961 | 0.948 |
| 61 | 0.000 (0.011); 0.947 | 0.000 (0.011); 0.947 | 0.000 (0.011); 0.948 | 0.000 (0.011); 0.956 | 0.946 |
| 40 | 0.000 (0.011); 0.954 | 0.000 (0.010); 0.952 | 0.000 (0.011); 0.955 | 0.000 (0.011); 0.957 | 0.946 |
| 25 | 0.000 (0.011); 0.947 | 0.000 (0.010); 0.941 | 0.000 (0.011); 0.949 | 0.000 (0.011); 0.942 | 0.949 |
| 10 | 0.000 (0.009); 0.952 | 0.000 (0.007); 0.928 | 0.000 (0.009); 0.958 | 0.000 (0.010); 0.836 | 0.958 |
| **No Heterogeneity, $\beta=0.05$** |  |  |  |  |  |
| 100 | 0.049 (0.011); 0.952 | 0.049 (0.011); 0.951 | 0.049 (0.011); 0.954 | 0.050 (0.011); 0.962 | 0.952 |
| 61 | 0.049 (0.011); 0.948 | 0.047 (0.011); 0.944 | 0.049 (0.011); 0.952 | 0.050 (0.011); 0.961 | 0.953 |
| 40 | 0.048 (0.011); 0.939 | 0.045 (0.011); 0.918 | 0.048 (0.011); 0.943 | 0.050 (0.012); 0.951 | 0.946 |
| 25 | 0.046 (0.011); 0.910 | 0.041 (0.010); 0.819 | 0.046 (0.011); 0.923 | 0.050 (0.012); 0.940 | 0.954 |
| 10 | 0.033 (0.010); 0.589 | 0.027 (0.008); 0.286 | 0.034 (0.011); 0.670 | 0.051 (0.012); 0.868 | 0.957 |
| **No Heterogeneity, $\beta=0.1$** |  |  |  |  |  |
| 100 | 0.099 (0.011); 0.945 | 0.097 (0.011); 0.945 | 0.099 (0.012); 0.950 | 0.100 (0.012); 0.963 | 0.946 |
| 61 | 0.098 (0.011); 0.932 | 0.095 (0.011); 0.920 | 0.098 (0.012); 0.944 | 0.100 (0.012); 0.956 | 0.947 |
| 40 | 0.097 (0.012); 0.911 | 0.091 (0.011); 0.859 | 0.097 (0.012); 0.933 | 0.100 (0.013); 0.954 | 0.951 |
| 25 | 0.092 (0.012); 0.844 | 0.083 (0.011); 0.649 | 0.092 (0.013); 0.896 | 0.101 (0.014); 0.947 | 0.955 |
| 10 | 0.065 (0.013); 0.348 | 0.055 (0.010); 0.094 | 0.072 (0.014); 0.518 | 0.102 (0.016); 0.895 | 0.964 |

Table 3 shows equivalent results when summary data sets of 25 SNPs are simulated under a multiplicative random effects model allowing for pleiotropy. The data are simulated so that the variability of the ratio estimates is twice that expected in the absence of pleiotropy (i.e the variance inflation parameter *ϕ* = 2). The performance of each approach follows a similar pattern to that presented for the fixed effect case in Table 2, with 1st order, 2nd order and iterative weights adversely affected by weak instrument bias and under coverage. The exact IVW estimate and its corresponding variance inflation parameter estimate are approximately unbiased. The non-parametric bootstrap procedure yields confidence intervals with approximately correct coverage. As before, confidence intervals have a tendency to be slightly conservative when the instruments are weak. Equivalent results for MR studies with 10 and 100 SNPs are shown in Appendix 6 of *Online Supplementary Material*. As the number of SNPs increases, the coverage of the exact IVW estimate’s confidence interval is increasingly closer to the nominal level.

**Table 3:** *Mean causal estimate 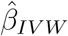, standard error (SE) and coverage frequency (CF) of the 95% confidence interval when using 1st order, 2nd order, iterative and exact weights. L=25. 2). 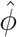 equals the variance inflation factor estimate (true value = 2)*.

| Mean | 1st order $w_j$ | 2nd order $w_j$ | Modified $w_j$ | | |
| --- | --- | --- | --- | --- | --- |
|  |  |  | Iterative | Exact |  |
| $F$ | $\hat{\beta}_{IVW}(SE); CF$ | $\hat{\beta}_{IVW}(SE); CF$ | $\hat{\beta}_{IVW}(SE); CF$ | $\hat{\beta}_{IVW}(SE); CF$ | $\hat{\phi}$ |
| **Heterogeneity, $\beta=0$** |  |  |  |  |  |
| 100 | 0.000(0.016); 0.949 | 0.000 (0.015); 0.950 | 0.000 (0.016); 0.950 | 0.000 (0.016); 0.939 | 2.000 |
| 61 | 0.000 (0.016); 0.950 | 0.000 (0.015); 0.951 | 0.000 (0.016); 0.951 | 0.000 (0.016); 0.940 | 2.004 |
| 40 | 0.000 (0.016); 0.953 | 0.000 (0.014); 0.951 | 0.000 (0.016); 0.955 | 0.000 (0.016); 0.944 | 1.999 |
| 25 | 0.000 (0.015); 0.949 | 0.000 (0.013); 0.945 | 0.000 (0.015); 0.954 | 0.000 (0.017); 0.945 | 2.003 |
| 10 | 0.000 (0.013); 0.952 | 0.000 (0.009); 0.924 | 0.000 (0.013); 0.960 | 0.000 (0.037); 0.970 | 1.943 |
| **Heterogeneity, $\beta=0.05$** |  |  |  |  |  |
| 100 | 0.050 (0.016); 0.948 | 0.048 (0.015); 0.947 | 0.050 (0.016); 0.949 | 0.050 (0.016); 0.938 | 2.002 |
| 62 | 0.049 (0.016); 0.951 | 0.046 (0.015); 0.943 | 0.049 (0.016); 0.954 | 0.050 (0.016); 0.943 | 1.998 |
| 40 | 0.048 (0.016); 0.949 | 0.044 (0.014); 0.924 | 0.048 (0.016); 0.953 | 0.050 (0.017); 0.943 | 1.995 |
| 25 | 0.046 (0.015); 0.933 | 0.039 (0.013); 0.839 | 0.046 (0.016); 0.940 | 0.051 (0.018); 0.944 | 1.987 |
| 10 | 0.033 (0.014); 0.719 | 0.025 (0.010); 0.378 | 0.034 (0.015); 0.778 | 0.051 (0.037); 0.960 | 1.967 |
| **Heterogeneity, $\beta=0.1$** |  |  |  |  |  |
| 100 | 0.099 (0.016); 0.947 | 0.096 (0.016); 0.942 | 0.099 (0.016); 0.952 | 0.100 (0.016); 0.942 | 2.005 |
| 61 | 0.098 (0.016); 0.941 | 0.092 (0.016); 0.922 | 0.098 (0.017); 0.951 | 0.100 (0.017); 0.941 | 2.004 |
| 40 | 0.097 (0.016); 0.932 | 0.088 (0.015); 0.862 | 0.097 (0.017); 0.947 | 0.101 (0.017); 0.940 | 2.003 |
| 25 | 0.092 (0.016); 0.888 | 0.078 (0.015); 0.676 | 0.093 (0.018); 0.924 | 0.101 (0.019); 0.942 | 2.003 |
| 10 | 0.065 (0.016); 0.456 | 0.051 (0.012); 0.131 | 0.072 (0.018); 0.639 | 0.101 (0.042); 0.956 | 2.023 |

### Power to detect a causal effect

In Appendix 7 of *Online Supplementary Material*, we show the power of 1st order, 2nd order, iterative and exact weighting to detect a causal effect for MR studies of 10, 25 and 100 SNPs when the data are generated from the same multiplicative random effects model. These simulations highlight a downside of exact weighting for causal estimation: when there are only a small number of weak instruments, its power can be considerably lower. For example, when *F* = 10 and the causal effect is 0.05 its power is just under half that of the 1st order IVW estimate (29% vs 13%). However, the power difference reduces considerably for 25 SNPs (e.g 60% vs 40%) and is effectively zero for 100 SNPs. The power of iterative weighting is much more comparable to that of 1st order weighting, but always slightly lower.

## Applied example

Figure 3 (top) shows a scatter plot of summary data estimates for the associations of 26 genetic variants with systolic blood pressure (SBP, the exposure) and coronary heart disease (CHD, the outcome). SNP-exposure association estimates were obtained from the International Consortium for Blood Pressure consortium (ICBP) [17]. SNP-CHD association odds ratios were collected from Coronary ARtery Disease Genome-Wide Replication And Meta-Analysis (CARDIoGRAM) consortium [18], which are plotted (and subsequently modelled) on the log odds ratio scale by making a normal approximation. These data have previously been used in a two-sample summary data MR analysis by Ference et al. [19] and Lawlor et al. [20], but we extend their original analysis here by applying our modified weights and conducting a more in depth inspection of each variant’s contribution to the overall heterogeneity. The mean *F*-statistic for these data is 61. Using 1st order weights the IVW estimate, which represents the causal effect of a 1mmHg increase in SBP on the log-odds ratio of CHD, is 0.053. This is shown as the slope of a solid black line passing through the origin. Cochran’s *Q* statistic based on 1st order weights is equal to 67.1, indicating the presence of substantial heterogeneity. For this reason, only random effects models were used to derive point estimates, confidence intervals and p-values for the causal effect.

**Figure 3:**
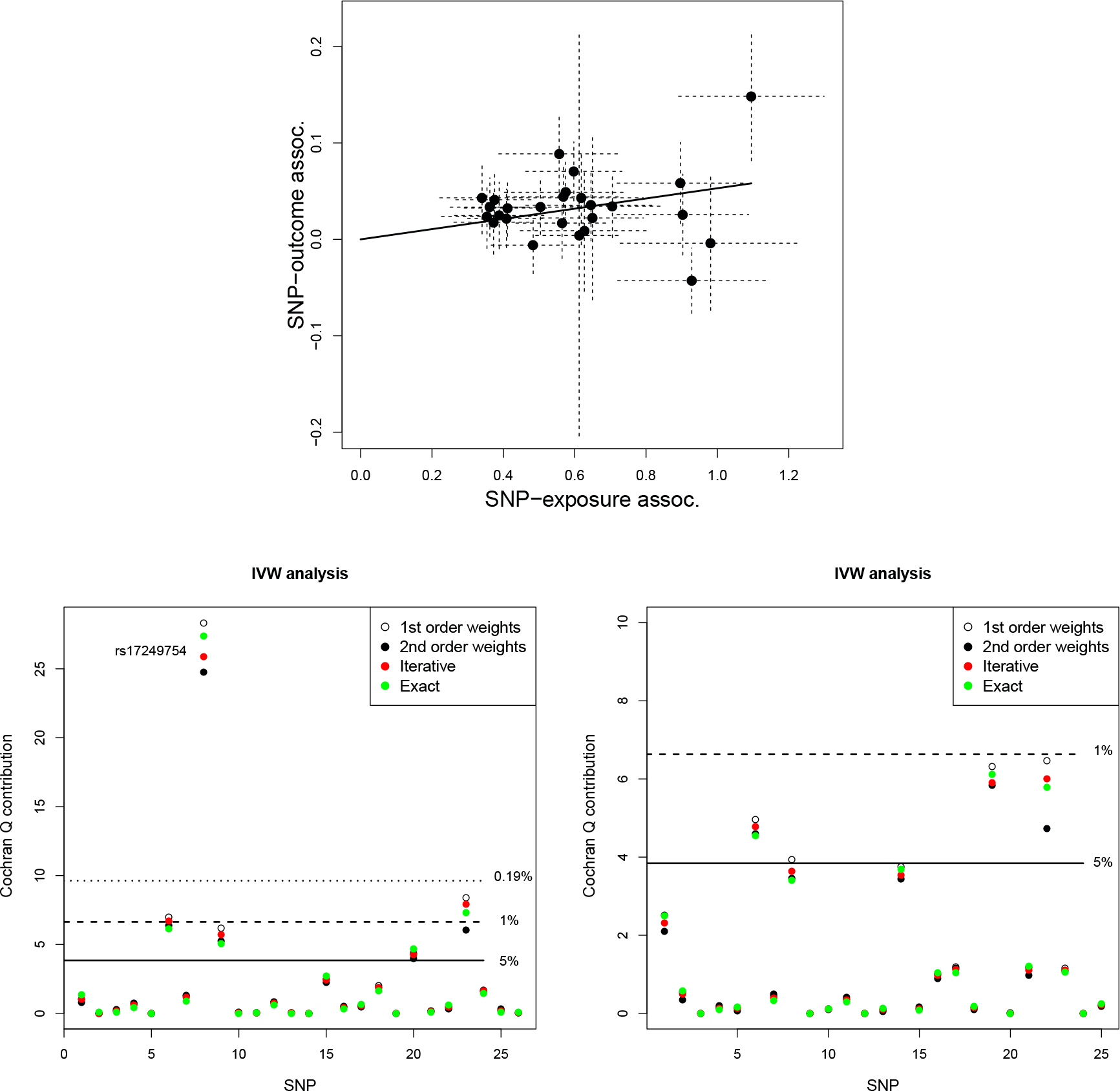
Top: Scatter plot of SNP-outcome associations 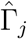 versus SNP-exposure associations 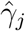. IVW estimate shown as a black slope. Bottom-left: Q contribution plots for the same data. Bottom-right: Q contributions after removal of rs17249754.

Table 4 shows the results of further IVW analyses using all weighting schemes. All schemes detect significant heterogeneity. As expected, the observed heterogeneity is largest when using 1st order weights, smallest when using 2nd order weights, and in between the two when using modified weights. Point estimates and standard errors are in good agreement across the different weights, because the mean instrument strength is high. Exact weighting gives the largest point estimate 0.054 under a random effects model. This is followed by 1st order and then 2nd order weights respectively. This ordering is as expected, given their relative susceptibility to regression dilution bias.

For comparison, we also report the Weighted Median [21], 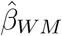, that can identify the causal effect when up to (but not including) half of the information in the analysis stems from genetic variants that are invalid IVs. Its estimate, which is calculated using 1st order weights, is 0.063. Although all approaches provide strong evidence in favour of a non-zero causal effect, the exact random effects IVW estimate is the least precise of all estimates. Consequently its p-value for testing the causal null hypothesis is the largest of all.

**Table 4:** *IVW and Weighted Median analyses of the causal effect of SBP on CHD risk for the complete data (top) and with SNP rs17249754 removed (bottom). 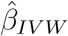 is the IVW estimate. 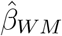 is the Weighted Median estimate. All IVW estimates fitted under a multiplicative random effects model (R.E), where 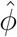 refers to the variance inflation factor estimate. The weighted median naturally accounts for heterogeneity via a bootstrapped variance.*

| Method (weights) | Estimate (C.I) | S.E. | P-value | Het. Stat (p) | $\hat{\phi}$ |
| --- | --- | --- | --- | --- | --- |
| **All 26 SNPs** |  |  |  |  |  |
| **Causal estimate** |  |  |  |  |  |
| IVW (1st -R.E) | $\hat{\beta}_{IVW}$: 0.053 (0.032,0.075) | 0.010 | $3.01 \times 10^{-5}$ | $Q = 67.1 (1.03 \times 10^{-5})$ | 2.68 |
| IVW (2nd -R.E) | $\hat{\beta}_{IVW}$: 0.050 (0.029,0.071) | 0.010 | $4.54 \times 10^{-5}$ | $Q = 58.8 (1.54 \times 10^{-4})$ | 2.35 |
| IVW (Iterative -R.E) | $\hat{\beta}_{IVW}$: 0.054 (0.032,0.075) | 0.010 | $2.40 \times 10^{-5}$ | $Q = 62.7 (4.43 \times 10^{-5})$ | 2.51 |
| IVW (Exact -R.E) | $\hat{\beta}_{IVW}$: 0.054 (0.027,0.082) | 0.014 | $4.60 \times 10^{-4}$ | $Q = 62.4 (4.84 \times 10^{-5})$ | 2.61 |
| **Weighted median (1st order weights)** |  |  |  |  |  |
| Weighted Median | $\hat{\beta}_{WM}$: 0.063 (0.042,0.084) | 0.011 | $4.90 \times 10^{-6}$ | - | - |
| **SNP rs17249754 removed** |  |  |  |  |  |
| **Causal estimate** |  |  |  |  |  |
| IVW (1st - RE) | $\hat{\beta}_{IVW}$: 0.066 (0.049,0.083) | 0.008 | $2.63 \times 10^{-8}$ | $Q = 35.0 (0.068)$ | 1.46 |
| IVW (2nd - RE) | $\hat{\beta}_{IVW}$: 0.063 (0.047,0.080) | 0.008 | $4.06 \times 10^{-8}$ | $Q = 30.6 (0.164)$ | 1.27 |
| IVW (Iterative - RE) | $\hat{\beta}_{IVW}$: 0.066 (0.049,0.083) | 0.008 | $2.90 \times 10^{-8}$ | $Q = 32.8 (0.107)$ | 1.37 |
| IVW (Exact - RE) | $\hat{\beta}_{IVW}$: 0.067 (0.049,0.085) | 0.009 | $8.37 \times 10^{-8}$ | $Q = 32.8 (0.108)$ | 1.39 |
| **Weighted median (1st order weights)** |  |  |  |  |  |
| Weighted Median | $\hat{\beta}_{WM}$: 0.065 (0.044,0.087) | 0.011 | $2.33 \times 10^{-6}$ | - | - |

Figure 3 (bottom-left) shows the individual contribution to Cochran’s *Q* statistic under each weighting scheme. Horizontal lines have been drawn to indicate the location of the 5th, 1st and 0.19th percentile of a 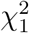 in order to help assess the magnitude of the contributions. The 0.19th percentile is derived as a 0.05 threshold adjusted for multiple testing using the Bonferroni correction. We see that the eighth SNP in our list (rs17249754) is responsible for the vast majority of the excess heterogeneity. Its contribution, *Q*_8_, ranges from approximately 24.5 to 28 depending on weighting. Variant rs17249754 sits in the ATPase plasma membrane Ca2+ transporting 1 (*ATP2B1*) gene, which is involved in intracellular calcium homeostasis, and is strongly associated with higher SBP. However, in the CARDIoGRAM consortium it is associated with reduced risk of CHD.

Since rs17249754 is also a strong instrument, and is potentially pleiotropic, its presence in the data could lead to the InSIDE assumption being violated. We therefore opt to remove it in a further sensitivity analysis, and Table 4 show the results. All IVW estimates increase by around 20% (lying between 0.063 and 0.067), but are ordered as before. Removal of rs17249754 leads to a dramatic reduction in the amount of heterogeneity present in the data, as referenced by *Q* statistics between 30 and 35 for all methods. Figure 3 (bottom-right) shows the updated contributions of each SNP to the various *Q* statistics after removing rs17249754. If only 1st order weighting were available, it might be tempting to exclude further variants from the analysis, but this signal is appropriately tempered when using exact weights. The Weighted Median estimate without rs17249754 is 0.065 (compared to 0.063 with). This highlights its inherent robustness to outliers, which is a major strength.

## Discussion

In this paper we have demonstrated the limitations of 1st and 2nd order weighting when used for IVW analysis in two-sample summary data Mendelian randomization. Most importantly, we highlight the potential for serious type I error inflation of Cochran’s *Q* statistic when using standard 1st order weights with weak instruments. In recent work, Verbanck et al. [22] also noted this same tendency and proposed a simulation-based alternative to 1st order weighting named ‘MR-PRESSO’. Our simulations show that modified weights can deliver much more reliable tests for heterogeneity than either 1st or 2nd order weighting, and offer a simple alternative to MR-PRESSO.

Modified weights were also shown to be a more reliable tool for the detection and removal of outliers in a given data set, as apposed to 1st order weights (which may detect too many outliers) and 2nd order weights (that may detect too few). Our simulations suggest that the exact weights should be used when testing for the overall presence of heterogeneity (referred to as the ‘global’ test by Verbanck et al. [22]) but that iterative weights are preferable if looking at the individual outliers. We suspect this is because exact weighting makes a more aggressive correction for regression dilution bias than iterative weighting. Its resulting estimate then makes more variants appear as outliers, because their ratio estimates are further away from the corrected slope. In effect, exact weighting leads to the detection of SNPs that are weak or pleiotropic.

An exciting finding of this paper is that the exact weighting also yields causal estimates that are remarkably robust to weak instrument bias. This opens up the potential for the significance threshold used to select SNPs as instruments to be set at a less stringent level. For example, in a specific analysis there might be four SNPs that are associated with the exposure with a p-value less than 5 × 10^−8^ (which equates to an F statistic of approximately 30 and above), but a total of 50 SNPs available that are associated with the exposure with a p-value less than 5 × 10^−6^ (which equates to an F statistic of approximately 20 and above). Modified weights would then be potentially preferable as a tool to more effectively utilize this larger set of SNPs within an MR analysis.

There are two downsides to the use of exact weights with weak instruments. Firstly, it can produce causal estimates with a reduced precision compared to simple 1st order weighting (although this difference disappears as the number of instruments increases). Secondly, if weak instruments are ‘discovered’ and analysed using the same data, then SNP-exposure estimates are more susceptible to the ‘winner’s curse’ than strong instruments. In preliminary work conducted in tandem with this paper, Zhao et al. [14] investigate the use of exact weighting for causal estimation and attempt to address both these issues. Specifically, they incorporate a penalized weight function within the exact weights. This reduces the effect of outliers (as apposed to explicit outlier removal) and increases the precision of the causal estimate. Sampling splitting is proposed to remove the affect of winner’s curse. The methods laid out in this paper differ from that of Zhao et al. [14] in four important ways. Firstly, we focus on the case of a multiplicative random effects pleiotropy commonly used in summary data MR whereas Zhao et al assume an additive random effects model. Secondly, Zhao et al derive and implement their method using profile-likelihood theory, whereas as our approach is motivated and implemented using Cochran’s *Q* statistic. Thirdly, we propose two forms of modified weighting (iterative and exact). Fourthly, we describe how both iterative and exact weighting can be used to test for heterogeneity as well as for causal estimation. For further details on the link between our work and that of Zhao et al. [14] see Appendix 1 of *Online Supplementary Material*.

### Limitations

Our conclusions regarding the use of modified weights are limited to the two-sample summary setting where SNP-outcome and SNP-exposure associations are estimated in independent but homogeneous samples. Further research would be required to decide if modified weights should be used in MR analyses of summary data estimates when there is partial overlap between samples, or in the single sample (total overlap) setting.

When Cochran’s *Q* statistic detects significant amounts of heterogeneity, it is prudent to test whether it is meaningfully biasing the analysis. This would indeed be the case if the heterogeneity were caused in part by directional pleiotropy with a non-zero mean. This would lead to bias in the IVW estimate, unless of course it was caused by a small number of SNPs that could be identified and removed from the analysis. MR-Egger regression [6, 7] could instead be used to address this. This approach simply regresses SNP-outcome associations on the SNP-exposure associations, tests for bias via its intercept, and estimates a bias-adjusted causal effect via its slope. Observed heterogeneity around the MR-Egger fit can then be quantified using an extended version of Cochran’s *Q* statistic, Ruücker’s *Q*′ [7, 23], and each variant’s contribution to *Q*′ can be used as the basis for outlier detection. Currently MR-Egger and Ruücker’s *Q*′ statistic use 1st order weights. Preliminary work suggests that modified weighting can be applied to MR-Egger regression to improve its performance - both in terms of causal effect estimation and heterogeneity quantification - just as for an IVW analysis, but further development and validation of this method is required.

Software to implement all of the methods introduced in this paper can be found within the `RadialMR` package to perform two-sample summary data MR, which can be downloaded from **https://github.com/WSpiller/RadialMR**.

##### Key messages

- Two-sample summary data Mendelian randomization requires the specification of inverse-variance weights for model fitting, heterogeneity quantification and outlier detection amongst a set of causal estimates.
- Heterogeneity indicates a possible violation of the necessary IV or modelling assumptions of which pleiotropy is a likely major cause.
- 1st order weights can inflate the type I error rate of Cochran’s *Q* statistic for detecting heterogeneity about the IVW estimate when the NOME assumption is strongly violated (as judged by a low *F*-statistic), and the true causal effect of interest is non-zero.
- 2nd order weights can reduce the power of Cochran’s *Q* statistic for detecting heterogeneity about the IVW estimate when the NOME assumption is violated.
- Modified weights (developed in this paper) preserve the type I error rate of Cochran’s *Q* statistic, whilst maintaining its statistical power.
- ‘Exact’ weights should be used for global tests of heterogeneity. ‘Iterative’ weights should be used to assess the outlier status of individual SNPs.
- IVW estimates obtained using exact weights are naturally corrected for regression dilution bias, and work well with large numbers of instruments, but can be imprecise relative to other weighting schemes with small numbers of weak instruments.
- Regardless of the number or strength of instruments used, 1st order weights always furnish unbiased IVW estimates and preserve the type I error rate under the causal null.

## Acknowledgements

We thank the reviewers and associate editor for providing helpful comments and suggestions that greatly improved this paper. Jack Bowden, Debbie A Lawlor and George Davey Smith all work in a Unit that receives support from the University of Bristol and UK Medical Research Council (MCUU00011/1, MCUU00011/2 and MCUU00011/6). The authors declare no conflict of interest.

